# Heterogeneous Responses of Hematopoietic Stem Cells to Inflammatory Stimuli are Altered with Age

**DOI:** 10.1101/163402

**Authors:** Mati Mann, Arnav Mehta, Carl de Boer, Monika S. Kowalczyk, Kevin Lee, Noga Rogel, Abigail R. Knecht, Daneyal Farouq, Aviv Regev, David Baltimore

**Affiliations:** Division of Biology and Biological Engineering, California Institute of Technology, Pasadena, CA 91125, USA.; David Geffen School of Medicine, UCLA, Los Angeles, CA 90025, USA.; Broad Institute of MIT and Harvard, Cambridge, MA 02142, USA.; Howard Hughes Medical Institute, Koch Institute of Integrative Cancer Biology, Department of Biology, Massachusetts Institute of Technology, Cambridge, Massachusetts 02140, USA

**Keywords:** Hematopoietic stem cells, stem cell aging, inflammation, single-cell RNA-sequencing

## Abstract

Long-term hematopoietic stem cells (LT-HSCs) maintain hematopoietic output throughout an animal's lifespan. With age, however, they produce a myeloid-biased output that may lead to poor immune responses to infectious challenge and the development of myeloid leukemias. Here, we show that young and aged LT-HSCs respond differently to inflammatory stress, such that aged LT-HSCs produce a cell-intrinsic, myeloid-biased expression program. Using single-cell RNA-seq, we identify a myeloid-biased subset within the LT-HSC population (mLT-HSCs) that is much more common amongst aged LT-HSCs and is uniquely primed to respond to acute inflammatory challenge. We predict several transcription factors to regulate differentially expressed genes between mLT-HSCs and other LT-HSC subsets. Among these, we show that *Klf5*, *Ikzf1* and *Stat3* play important roles in age-related inflammatory myeloid bias. These factors may regulate myeloid versus lymphoid balance with age, and can potentially mitigate the long-term deleterious effects of inflammation that lead to hematopoietic pathologies.

**Highlights:**

- LT-HSCs from young and aged mice have differential responses to acute inflammatory challenge.
- HSPCs directly sense inflammatory stimuli *in vitro* and have a robust transcriptional response.
- Aged LT-HSCs demonstrate a cell-intrinsic myeloid bias during inflammatory challenge.
- Single-cell RNA-seq unmasked the existence of two subsets within the LT-HSC population that was apparent upon stimulation but not steady-state. One of the LT-HSC subsets is more prevalent in young and the other in aged mice.
- *Klf5*, *Ikzf1* and *Stat3* regulate age‐ and inflammation-related LT-HSC myeloid-bias.

**One sentence summary:** Murine hematopoietic stem cells display transcriptional heterogeneity that is quantitatively altered with age and leads to the age-dependent myeloid bias evident after inflammatory challenge.

## Introduction

Long-term hematopoietic stem cells (LT-HSCs) must overcome the stresses of aging to maintain appropriate immune cell output throughout a human's life (Akunuru and Geiger, 2016; Chen et al., 2016; Denkinger et al., 2015; Dykstra et al., 2011; Geiger et al., 2013; Mehta et al., 2015; Morita et al., 2010; Sawai et al., 2016). These stresses include replicative stress (Bernitz et al., 2016; Flach et al., 2014; Wang et al., 2012), as well as acute and chronic infectious challenge (King and Goodell, 2011; Nagai et al., 2006). Hematopoietic stem and progenitor cells (HSPCs) express innate immune receptors (King and Goodell, 2011), such as toll-like receptors (TLRs), and respond to many inflammatory mediators, including IFN-γ (Baldridge et al., 2010), M-CSF (Mossadegh-Keller et al., 2013), and the gram-negative bacterial component lipopolysaccharide (LPS) (Nagai et al., 2006). In response to acute LPS exposure, LT-HSCs increase proliferation, mobilize to the peripheral bloodstream (King and Goodell, 2011), and initiate emergency myelopoiesis to increase the system's output of innate immune cells (Haas et al., 2015). This increased output may be mediated by hematopoietic progenitors, such as multipotent progenitors (MPPs) (Pietras et al., 2015; Young et al., 2016), in part due to direct secretion of cytokines that drive myeloid differentiation (Zhao et al., 2014).

Physiologic aging in both humans and mice leads to permanent changes in LT-HSC function, such as myeloid-biased hematopoietic output (Akunuru and Geiger, 2016). This is often accelerated by chronic inflammation and, when dysregulated, can lead to replicative exhaustion and extramedullary hematopoiesis (Esplin et al., 2011; Mehta et al., 2015). Several hypotheses have been proposed to explain the age related changes in LT-HSC function (Kovtonyuk et al., 2016). First, cell-intrinsic changes within each aged LT-HSC might make it inherently myeloid-biased (Grover et al., 2016). Second, the LT-HSC population may be comprised of subsets of myeloid‐ and lymphoid-biased cells, the composition of which changes with age such that myeloid-biased LT-HSCs are more prevalent within the aged LT-HSC population (Dykstra et al., 2007; Yamamoto et al., 2013). The true nature of these age-related changes may in fact be a combination of both of these hypotheses, such that with age there is a growing subset of more intrinsically myeloid-biased LT-HSCs.

The transcriptional state of LT-HSCs in steady state and in response to inflammatory mediators may help shed light on these questions, but is currently still poorly understood. A number of epigenomic and transcriptomic changes have been observed during bulk and single-cell analysis of young and aged LT-HSCs (Cabezas-Wallscheid et al., 2014; Grover et al., 2016; Kowalczyk et al., 2015; Mehta and Baltimore, 2016; Sanjuan-Pla et al., 2013; Sun et al., 2014; Yu et al., 2016). However, it is unclear if and how these changes lead to altered LT-HSC function, as seen with age-related myeloid bias (Dykstra et al., 2011; Gekas and Graf, 2013). In particular, a recent study using single-cell RNA-seq (scRNA-seq) (Kowalczyk et al., 2015) of steady-state, resting LT-HSCs has not identified a subpopulation structure; however, an appreciation of the cell-intrinsic differences between young and aged LT-HSCs may become apparent in the setting of acute stress, such as when analyzing their response to inflammatory challenge. An understanding of how the response of LT-HSCs to inflammatory mediators changes with age may therefore help elucidate the underlying mechanism of age-related myeloid bias. This may further provide insight into age-related pathologies such as improper immune responses to vaccines or infectious challenge, and the development of myeloid leukemia.

In this work, we investigate the acute inflammatory response of mouse HSPCs *in vitro* and *in vivo*, and how this response may be altered with age (Figure 1A). We demonstrate that major HSPC subtypes respond transcriptionally to inflammatory stimuli and that this response is similar to the prototypical response of mouse bone marrow derived dendritic cells (BMDCs). Using *in-vivo* experiments, we show that the age-dependent myeloid bias after inflammatory challenge is intrinsic to LT-HSCs. Using single-cell RNA-seq (scRNA-seq) we find that the LT-HSC compartment is comprised of at least two subsets that become apparent only upon stimulation. One of these subsets has features consistent with myeloid-bias, with distinct cell-intrinsic responses to inflammatory stimulation. The myeloid-biased subset increases dramatically with age. We identify putative transcriptional regulators of these cell states, and demonstrate the role of these regulators in age-related myeloid bias and differential responses to TLR ligands.

**Figure 1.**
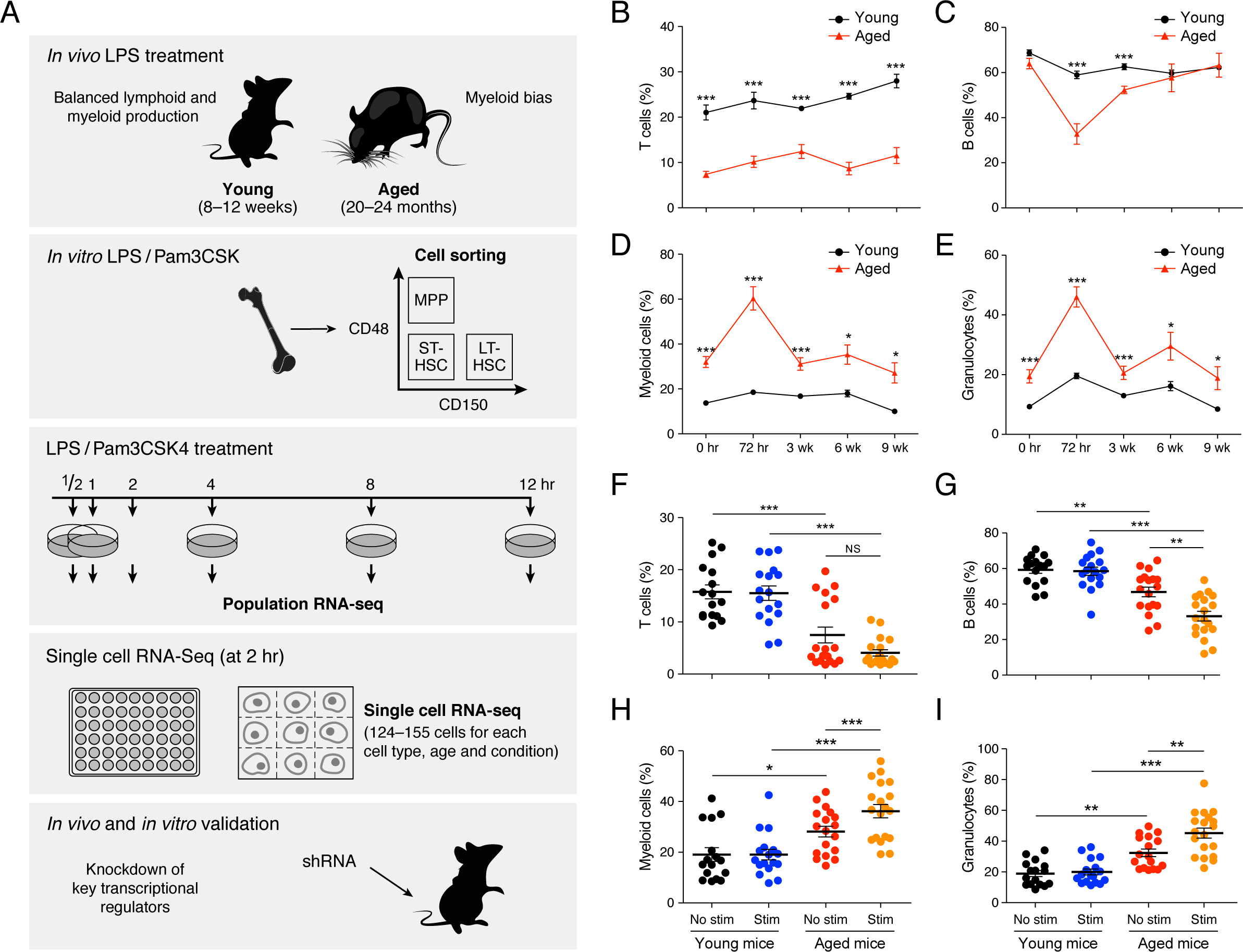
Aged hematopoietic stem cells exposed to inflammatory signals demonstrate increased myeloid output in a cell-intrinsic manner. (A) Schematic overview of the approach. (B)-(C) Young (8-12 weeks) and aged (20-24 months) mice were exposed to single sub-lethal dose of LPS and peripheral blood (B) T cell, (C) B cell, (D) myeloid cell and (E) granulocyte frequencies were measured by flow cytometry at the indicated time points after LPS exposure (n = 4-14 per group). (F)-(I) LT-HSCs (LincKit+Sca1+CD150+CD48-) sorted from young and aged CD45.2 mice were stimulated with LPS and Pam3csk4 for 2 hours prior to competitive transplant into CD45.1 recipients. Peripheral blood (F) CD3+, (G) CD19+, (H) CD11b+ and (I) Gr-1+ frequencies were measured by flow cytometry at 3 months post-reconstitution (n = 11-12 per group). Data represent at least two independent experiments and are presented as mean ± SEM. * denotes p < 0.05, ** denotes p < 0.01 and *** denotes p < 0.001. P-values were corrected for multiple hypothesis testing by Bonferroni's multiple comparison test.

## Results

### Differential response of young and aged mice to LPS in vivo

To investigate the acute inflammatory response of hematopoietic progenitors from mice at different ages, we challenged 8-12 week old (‘young’) and 20-24 month old (‘aged’) mice with a single intraperitoneal injection of LPS (0.5*µ*g/kg, Figure 1B-E). Young mice responded with a gradual increase in the frequency of peripheral blood T cells over 72 hours, which was sustained 9 weeks after the LPS challenge (Figure 1B). Conversely, aged mice had an over twofold lower baseline frequency of T cells in their peripheral blood compared to young mice, and the frequency of these cells remained unchanged after the LPS challenge (Figure 1B). Both young and aged mice showed a decrease in peripheral blood B cell frequencies after LPS treatment. However, the acute response was particularly dramatic in aged mice, which had a twofold loss in the frequency of B cells by 72 hours, and then recovered to levels comparable to those seen in young mice by 6 weeks post-challenge (Figure 1C). In contrast to the milder increase in the myeloid output of young mice, there was an over twofold increase in peripheral blood myeloid frequencies in aged mice by 72 hours post-challenge (Figure 1D,E). This eventually normalized to baseline frequencies by 9 weeks post-challenge. Aged mice therefore demonstrated a strong acute increase in myeloid output in response to inflammatory challenge; this was not observed in young mice, which conversely responded primarily with an increase in T cell output.

To evaluate the cumulative effect of acute inflammatory challenges on myeloid output, we performed an LPS boost of all cohorts 2 months after the initial challenge. This resulted in a dramatic upregulation of peripheral blood myeloid cells in aged mice lasting at least 12 days, whereas again, only a milder increase in myeloid output was seen in young mice (**Figure S1A**). The spleens of stimulated aged mice revealed increased myeloid cell frequencies and a dramatic loss of T cells compared to young mice (**Figure S1B**). Finally, bone marrow of stimulated aged mice had a three-fold enrichment for LT-HSCs compared to stimulated young mice, with a milder enrichment in short-term HSCs (ST-HSCs) and MPPs (**Figure S1C**). This enrichment in LT-HSCs is higher than the two-fold enrichment seen between unstimulated aged and young mice (Beerman et al., 2010; Mehta et al., 2015), suggesting that acute inflammatory stimuli promote myeloid-biased output from HSPCs and that the differences in immune cell output might originate from LT-HSCs.

### Aged LT-HSCs stimulated with inflammatory signals yield a distinctive, long-term myeloid-biased output

To examine whether the age-related myeloid bias in hematopoietic output after inflammatory challenge is intrinsic to LT-HSCs, we tested the impact of stimulation on the ability of young and aged LT-HSCs to reconstitute the immune system. Specifically, we first sorted LT-HSCs, ST-HSCs and MPPs from young and aged CD45.2 C57BL/6 mice (**Figure S2**). The cells were then either maintained unstimulated or stimulated with LPS and Pam3csk4 for 2 hours *in vitro*. They were subsequently transplanted together with CD45.1 bone marrow helper cells into lethally irradiated young CD45.1 C57BL/6 mice. Peripheral blood counts of CD45.2 expressing cells were monitored for several months (Figure 1F-I and **Figure S3A,B**). Both unstimulated and stimulated, young and aged LT-HSCs demonstrated long-term reconstitution of the immune system in these primary transplants. All transplanted ST-HSCs maintained hematopoietic output for up to 3 months, whereas all MPPs failed long-term reconstitution (**Figure S3A,B**), as expected. To validate which of these cells maintain long-term reconstitution potential, we performed secondary transplants by taking bone marrow donor cells from primary transplant mice and transplanting them to lethally irradiated young CD45.1 mice. Bone marrow from mice initially transplanted with CD45.2 LT-HSCs successfully reconstituted the immune system with CD45.2 immune cells while bone marrow from mice transplanted with CD45.2 ST-HSCs or MPPs failed to do so (**Figure S3C,D**).

Notably, stimulated aged LT-HSCs led to a distinctive immune cell output in the reconstitution experiments, with a particularly marked myeloid bias compared to all other conditions (Figure 1F-I). At 3 months post-reconstitution (*i.e.* 3 months after the *in vitro* LPS/Pam3csk4 challenge), no difference in the frequencies of peripheral blood myeloid and lymphoid cells were seen between mice reconstituted with either stimulated or unstimulated young LT-HSCs (Figure 1F-I). As previously reported (Beerman et al., 2010; Pang et al., 2011), unstimulated aged LT-HSCs had higher peripheral blood myeloid output and lower lymphoid output compared to unstimulated young LT-HSCs (Figure 1F-I). However, stimulated aged LT-HSCs demonstrated a further decrease in the frequency of peripheral blood lymphoid cells (Figure 1F,G) and a marked increase in the frequency of peripheral blood myeloid cells (**Figure H,I**), even compared to the unstimulated aged LT-HSCs. Thus, aged LT-HSCs demonstrated myeloid-biased 'memory’ of the initial *in vitro* LPS/Pam3csk4 challenge that persisted for several months post-reconstitution, a phenomenon not seen with stimulated young LT-HSCs. Interestingly, no significant difference in LT-HSC frequency, including the previously identified myeloid-biased CD41+ LT-HSC subpopulation (Gekas and Graf, 2013), was observed between cohorts (**Figure S3E-H**).

### HSPCs demonstrate a canonical transcriptional response to TLR ligands

We hypothesized that the differential effects of young and aged stimulated LT-HSCs may be due to a variable immediate transcriptional response to inflammatory signals. To test this hypothesis, we measured the transcriptional profiles of populations of HSPCs from young and aged mice during a 12-hour time-course of LPS/Pam3csk4 stimulation *in vitro* (Figure 2A). LT-HSCs, ST-HSCs and MPPs from both young and aged mice all demonstrated a robust and similar transcriptional response (Figure 2B), which largely resembled that seen in mature cell types with different physiological functions, such as BMDCs after LPS stimulation (Figure 2B-D) (Jovanovic et al., 2015). This includes the same temporal ordering of induction in inflammatory gene clusters (Figure 2C) as in mature cell types (Amit et al., 2009; Ramirez-Carrozzi et al., 2009), up-regulation of NF-KB-related genes (Figure 2C) (Bhatt et al., 2012; Hao and Baltimore, 2009), and induction of the expression of several effector cytokines (Figure 2D), albeit at slightly lower levels (**Figure S4A-C**). Thus, the response of young and aged HSPCs to inflammatory activation follows the canonical response of mature cells to similar stimulation, both in the identity of the regulated genes and in the timescale of the response. This suggests that the differences in the reconstitution outcome could not be resolved by the differences in the transcriptional response when measured at the population level.

**Figure 2.**
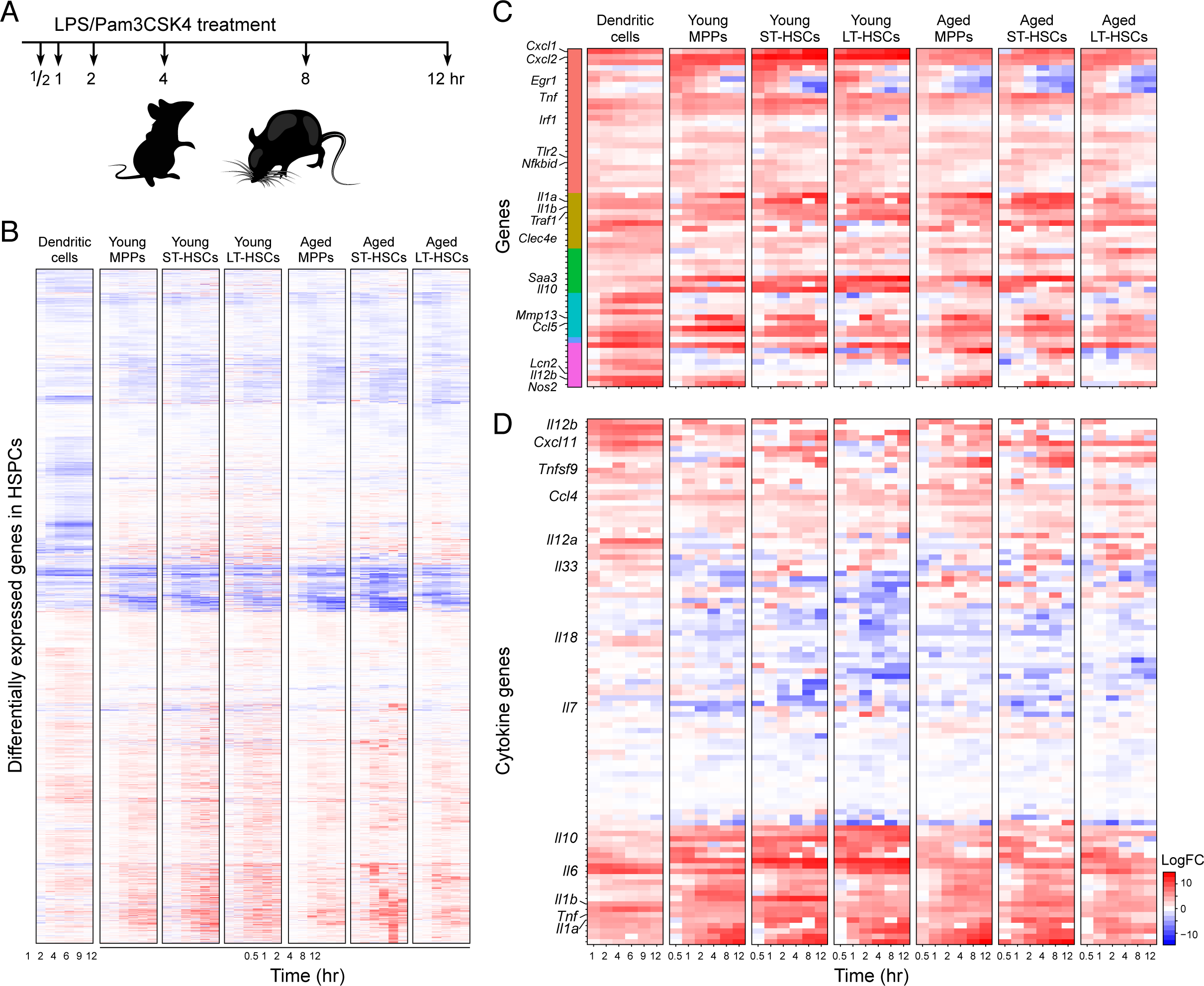
Early hematopoietic progenitors demonstrate a rapid transcriptional response to inflammatory signals. (A) Schematic of LPS and Pam3CSK4 time-course experiment. LT-HSCs, ST-HSCs and MPPs from young (8-12 weeks) and aged (20-24 months) mice were exposed to LPS and Pam3CSK4 *in vitro* for the indicated time after which RNA was harvested for bulk RNA-sequencing. (B) Heatmap of differentially expressed genes in young and aged hematopoietic progenitors alongside an expression map of mature bone-marrow derived dendritic cells (DCs) challenged with LPS for comparison. (C) Heatmap of NF-kB regulated inflammatory genes clustered by temporal expression patterns described previously (Bhatt et al., 2012). (D) Heatmap of cytokines expressed in early progenitors.

### Single-cell RNA-seq reveals two subsets of LT-HSCs with distinct responses to stimulation and compositional changes with age

Next, we considered the possibility that there are different subsets of LT-HSCs either in steady state or post-stimulation (“cell intrinsic changes”), and that their relative proportions may change with age (“compositional changes”). Early hematopoietic progenitors are comprised of heterogeneous functional subpopulations (Benz et al., 2012; Gekas and Graf, 2013; Morita et al., 2010; Sanjuan-Pla et al., 2013), which often reveal themselves in response to inflammatory stimuli (Haas et al., 2015; Zhao et al., 2014). While a previous scRNA-seq study of LT-HSCs has mostly revealed age-related differences in the cell cycle (Kowalczyk et al., 2015), we hypothesized that stimulation could unveil additional cell intrinsic distinctions that cannot be observed in resting cells.

To determine the composition of HSPCs in each age group and condition, we performed full-length scRNA-seq (Picelli et al., 2013) of young and aged LT-HSCs, ST-HSCs and MPPs, with and without 2 hours of *in vitro* LPS/Pam3csk4 stimulation. As we aimed to distinguish cell intrinsic, possibly subtle, stimulus-specific states within a very well-defined cell population, we opted for the deeper-coverage full length scRNA-Seq approach over massively parallel approaches (Tanay and Regev, 2017; Wagner et al., 2016). We profiled 2046 individual cells from nine mice (5 young, 4 aged), with 124-186 cells for each given cell type and condition. In order to eliminate sources of variability resulting from known confounding factors, we removed 611 cells as low-quality and 58 as possible contaminants (**STAR-Methods**). In addition, 578 of the cells were actively cycling (**STAR-Methods**). Overall, we retained 949 cells for subsequent analysis, comprised of 187 MPPs, 404 ST-HSCs, and 358 LT-HSCs.

We identified three major groups of cells using unsupervised clustering (see **STAR-Methods**) (Figure 3A; denoted clusters 1, 2 and 3). Cluster 1 (311 cells) contained most (302 of 354) of the unstimulated HSPCs of all types, forming a continuum from MPPs to LT-HSCs (Figure 3B), with aged and young LT-HSCs clustering together (Figure 3D), and hardly any stimulated cells. Clusters 2 (103 cells) and 3 (421 cells) almost exclusively contained stimulated HSPCs (Figure 3C), and had opposing patterns with respect to aged and young LT-HSCs (Figure 3E): cluster 3 contained 77% of the aged stimulated LT-HSCs and only 13% of the young stimulated LT-HSCs (Figure 3E), whereas cluster 2 had 72% of the young stimulated LT-HSCs, and only 10% of the aged stimulated LT-HSCs (Figure 3E). This suggests that there are distinct subsets of LT-HSCs in the bone marrow that can be discerned by their different cell intrinsic responses to stimulation, and that the relative frequencies of these subsets appear to change with age. Of note, the distinction between these LT-HSC subsets could only be discerned with stimulation. Given these findings, we focused further analysis on LT-HSCs.

**Figure 3.**
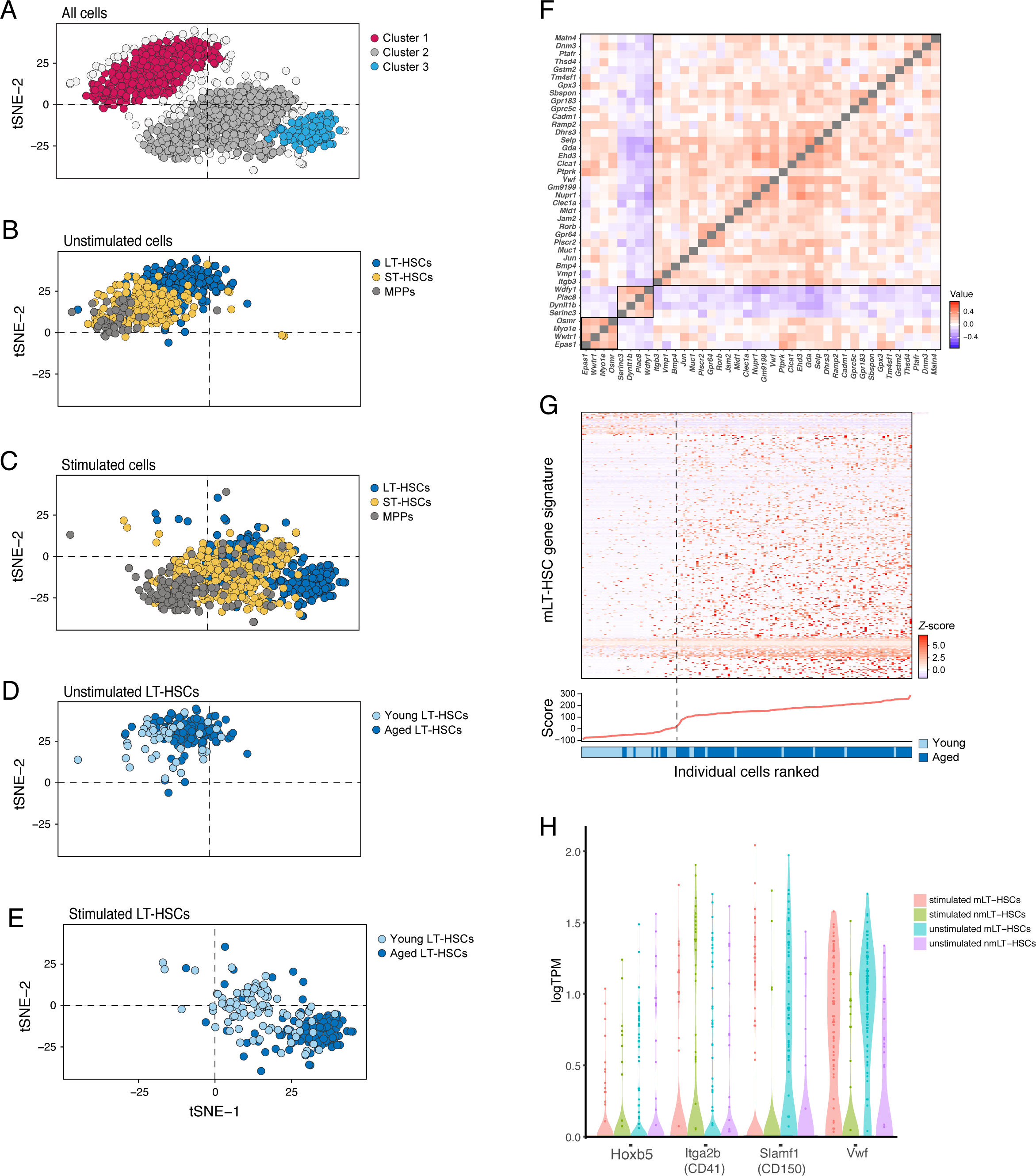
Single-cell RNA-sequencing reveals heterogeneity in the hematopoietic stem cell compartment that is altered with age. (A)-(E) LT-HSCs, ST-HSCs and MPPs from young and aged mice were stimulated with LPS and Pam3CSK4 and sorted for single-cell RNA-seq. Single cell RNA-seq data for all cells, projected on two t-SNE axes. (A) Density-based clusters. (B)-(E) Single-cell t-SNE plots (as before) indicating all cell types among (B) unstimulated and (C) stimulated cells, and mouse ages among (D) unstimulated and (E) stimulated cells. (F) Correlation across cells between DE genes common to both simulated cluster 3 vs 2 and unstimulated aged vs young LT-HSCs. Three clusters of correlated genes are identified. (G) Heat-map represents the expression values of genes in the unstimulated myeloid-biased gene signature for each single unstimulated LT-HSC. The panels below show the myeloid signature score for each cell and is the basis for the ordering of the x axis. The color coded bar at the bottom shows the age of the animals from which the cells were derived. (H) Violin plots of all LT-HSCs showing the mRNA expression of *Hoxb5*, *CD41*, *CD150* and *Vwf*.

### Stimulated aged LT-HSCs express a myeloid-biased gene expression signature

To identify the differences between the stimulated LT-HSCs in cluster 3 and cluster 2, we examined genes that were differentially expressed between the two clusters. Cluster 3-specific genes were enriched for genes related to myeloid function and inhibiting lymphoid differentiation, including pathways related to NF-κB localization, negative regulation of lymphocyte development, macrophage proliferation, cell migration and localization, and platelet derived growth factor signaling (**Figure S4D**). Cluster 2-specific genes were enriched for genes involved in lymphocyte development, cell proliferation, and the acute inflammatory response (**Figure S4E**). Of note, cells in any given cluster, regardless of age, were similar to each other and distinctly different from cells in the other cluster. Interestingly, we observed some differences in the level of expression of inflammatory genes such as IL6 and TNFα in stimulated LT-HSCs in both clusters (**Figure S4F,G**). This data suggests that aged and young LT-HSCs have different proportions of cells that display unique lineage-biased pathway preferences in response to inflammatory signals.

### Myeloid-biased LT-HSCs can be identified by a distinct signature in unstimulated LT-HSCs and their proportion increases with age

To test whether the LT-HSC subsets exist also in unstimulated cells in steady-state conditions, we identified the 47 genes that were differentially expressed both when comparing cluster 3 *vs.* cluster 2 (SCDE FDR < 0.01) and when comparing unstimulated aged *vs.* young LT-HSCs within cluster 1 (**Figure S5A; STAR-Methods**, SCDE FDR < 0.1). We then tested whether these 47 genes coherently co-vary across the 149 unstimulated LT-HSCs, and thus might reflect a variable cell state within these cells. Indeed, we identified three distinct co-varying gene clusters (FIgure 3F), two of which contained genes involved in myeloid and platelet differentiation, including *Selp*, *Vwf*, *Gpr64*, *Plscr2*, and *Wdfy1*. Notably, recent studies have reported myeloid-biased CD41, Vwf or CD150-high expressing LT-HSC subpopulations (Dykstra et al., 2011; Gekas and Graf, 2013; Sanjuan-Pla et al., 2013); in our analysis, aged LT-HSCs have increased yet variable expression of CD150 and Vwf, and to a lesser extent expression of CD41 (Figure 3H **and Figure S4H-J**). A recent report has also suggested Hoxb5 can be used as a marker for truly long-term reconstituting subsets of LT-HSCs (Chen et al., 2016); our results show no significant difference in Hoxb5 expression among the various subsets (Figure 3H).

Next, we generated a refined gene signature by first scoring unstimulated LT-HSCs with the initial set of 47 genes (**Figure S5A**), identifying two putative cell subsets (**STAR Methods**). We then used these subsets to initialize *k*-means clustering (k=2) within the unstimulated LT-HSCs. We used the identities of the cells based on this clustering to designate them as myeloid-biased LT-HSCs, or “mLT-HSCs”, and non-myeloid biased LT-HSCs, or “nmLT-HSCs”. We next tested these two final clusters for differentially expressed genes, finding 365 upregulated genes and 34 downregulated genes in the mLT-HSC cluster, which we use to define our “mLT-HSC signature” (SCDE FDR < 0.1 and the same direction of change as for stimulated mLT-HSCs) (**Table S1**).

In the final *k*-means clusters, 92% of cells in the myeloid biased cluster were aged cells and only 8% were young cells (Figure 3G, to the right of the dashed line), while only 20% of cells in the non-myeloid biased cluster were aged cells and 80% were young cells (Figure 3G, to the left of the dashed line). This is consistent with our findings that the frequency of stimulated mLT-HSCs increases with age (Figure 3E). Applying the same signature to our stimulated LT-HSCs or to an independent dataset of unstimulated aged and young LT-HSCs from two mouse strains (Kowalczyk et al., 2015) showed consistent results: while the LT-HSC population is inherently heterogenous, more aged LT-HSCs score highly for the mLT-HSC signature (**Figure S5B,C**). Thus, the mLT-HSC signature allowed us to identify the subtle portion of myeloid-biased like cells among the young unstimulated LT-HSCs, and to show that the proportion of high-scoring mLT-HSC cells rises with age.

### Identification of transcription factors regulating LT-HSC subpopulations

To identify transcription factors (TFs) that may regulate differentially expressed genes between the mLT-HSC and nmLT-HSC subsets, we looked for enriched TF motifs in the enhancer sequences associated with these genes (Lara-Astiaso et al., 2014) (Figure 4A,B). In particular, we focused on TFs which themselves were differentially expressed between mLT-HSCs and nmLT-HSCs (Figure 4A,B), since altered transcriptional regulation of a TF is likely to affect its target genes.

**Figure 4.**
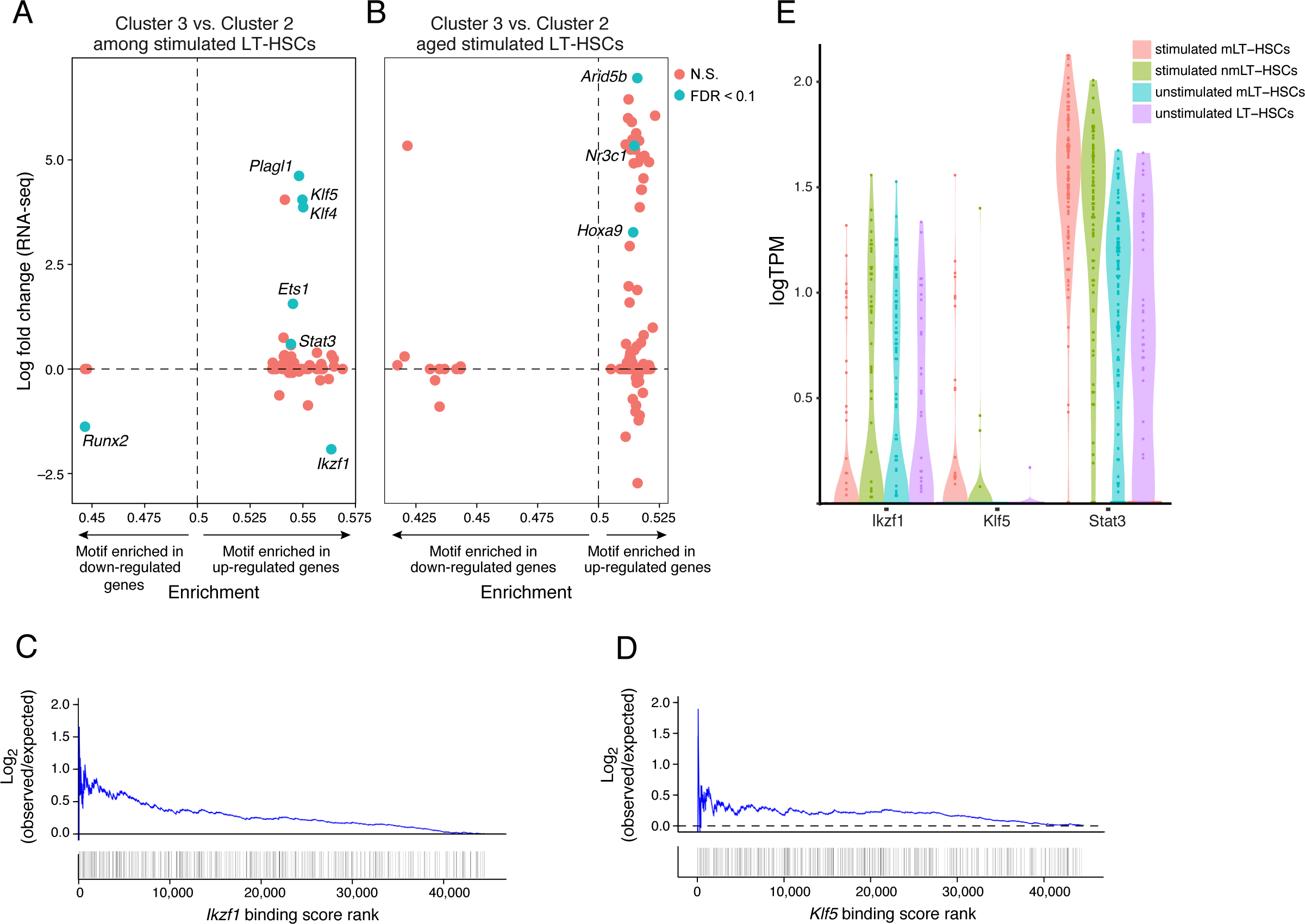
Several transcriptional factors may be involved in the underlying myeloid-bias of aged LT-HSCs. (A) Enrichment of transcription factor motifs in enhancers of cluster 3-vs-2-specific genes (x-axis) and differential expression of the TF genes themselves in the same comparison (y-axis). Significant genes (FDR<0.1) are indicated. An enrichment score > 0.5 indicates that the TF motif is enriched among genes that are expressed more highly in cluster 3, while a score below 0.5 indicates that the TF is expressed more highly in cluster 2. (B) As in (A), but considering only aged LT-HSCs in cluster 3 and cluster 2. (C,D) Observed/expected enrichment of enhancers associated with DE genes (indicated by heatmap), when sorted by decreasing motif strength for (C) Ikzf1 and (D) Klf5. (E) Violin plots of all LT-HSCs showing the mRNA expression of *Ikzf1*, *Klf5* and *Stat3*.

Among the 10 significant TFs (Figure 4A,B; blue dots) were *HoxA9*, *Klf4*, *Klf5*, *Ikzf1*, and *Stat3* (Figure 4C-E **and Figure S5D-F**). *HoxA9* is known to potentiate LT-HSC function (Lebert-Ghali et al., 2016) and lead to LT-HSC proliferation (Lebert-Ghali et al., 2016; Smith et al., 2011); interestingly, it was upregulated in the aged stimulated mLT-HSCs (comparing aged LT-HSCs between cluster 3 and 2; Figure 4B). The role of *Klf4*, *Klf5* and *Ikzf1* in LT-HSC function is less well established, although it has been suggested that *Klf5* may play a role in LT-HSC homing to the bone marrow niche (Taniguchi Ishikawa et al., 2013). All four TFs had their motifs enriched in the enhancers of genes upregulated in mLT-HSCs (Figures 4A-D). Among these, *Klf4* and *Klf5* were transcriptionally upregulated in mLT-HSCs, whereas *Ikzf1* was down-regulated in mLT-HSCs, and had motif instances enriched in genes upregulated in these cells (Figure 4A,E), consistent with its known role as a repressor (Koipally et al., 1999). *Stat3* is a known regulator of HSC self-renewal, especially under stress conditions, (Chung et al., 2006) and loss of *Stat3* is associated with a LT-HSC aging phenotype (Mantel et al., 2012). *Stat3* binding sites were enriched at enhancers of genes upregulated in mLT-HSCs (Figures 4A,E).

### Klf5, Ikzf1 and Stat3 play a role in age-related inflammatory myeloid bias of LT-HSCs

We tested the predicted role of these TFs in the age-related myeloid bias of LT-HSCs by shRNA knockdowns of each of *HoxA9*, *Klf4*, *Klf5*, *Ikzf1* or *Stat3* in young and aged HSPCs, leading to a 50-75% reduction in mRNA expression of each gene (**Figure S6A**). We compared these knockdowns to a control empty vector (MG) and knockdown of *Zbtb4*, a gene that is expressed in LT-HSCs but not differentially expressed between mLT-HSCs and nmLT-HSCs. HSPCs were first isolated from young and aged mice, and subsequently transduced with a retroviral vector expressing an shRNA construct for a particular TF. These cells were then used to reconstitute lethally irradiated young C57BL/6 recipient mice. As expected, at 3-months post reconstitution, when all the cells produced are progeny of transplanted LT-HSCs, we found that mice reconstituted with control (MG vector) young LT-HSCs had higher lymphoid output (Figure 5A, compare black bars to dark red bars), and lower myeloid (CD11b+) and granulocyte (Gr-1+) output (Figure 5B,C, compare black bars to dark red bars) than mice reconstituted with control aged LT-HSCs.

**Figure 5.**
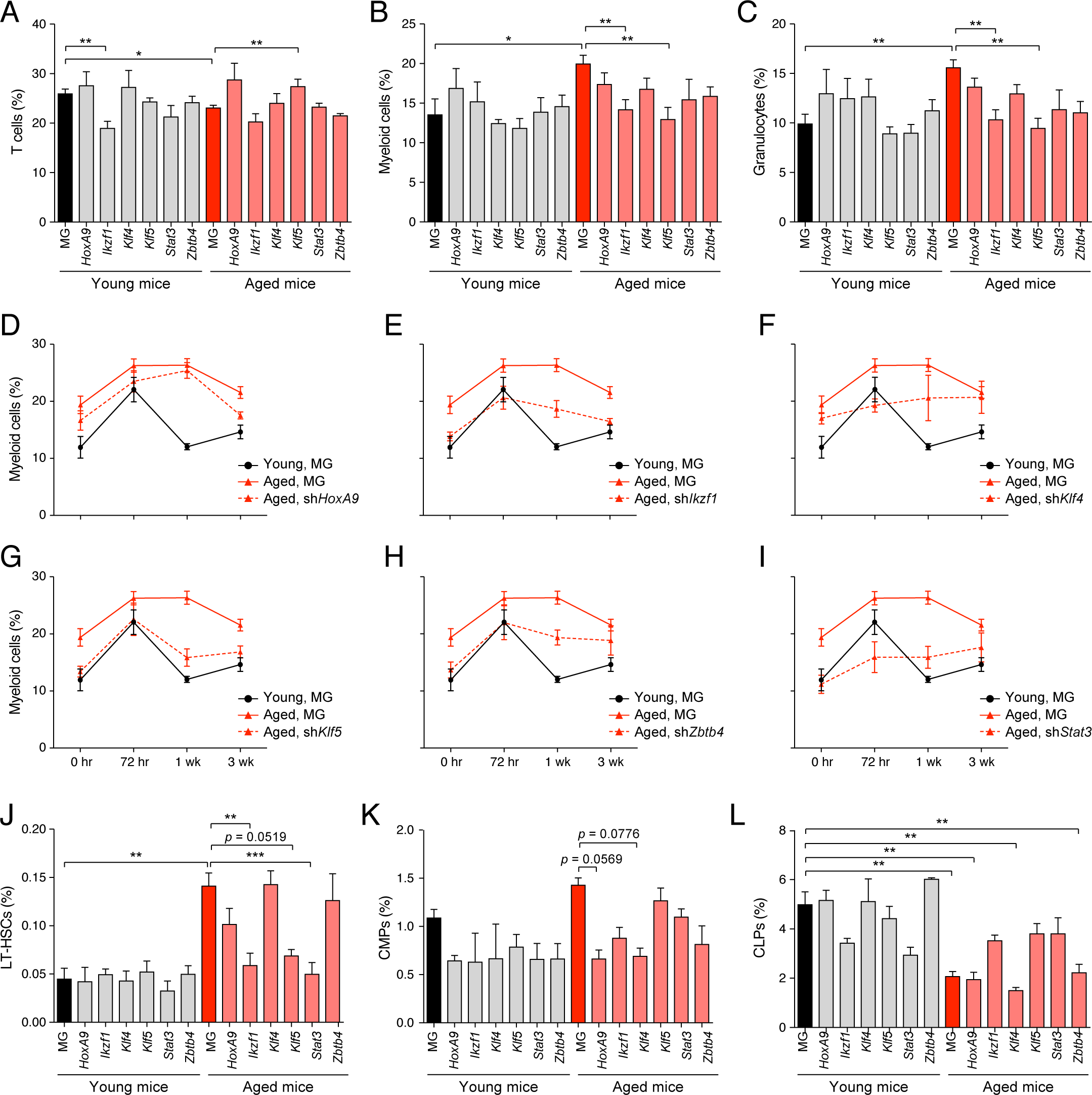
Klf5, Ikzf1, and Stat3 regulate steady-state and inflammatory age-related myeloid bias. (A)-(L) Bone marrow cells from young (8-12 weeks) and aged (20-24 months) C57BL/6 mice were transduced with constructs to knock-down the indicated transcription factors. These cells were subsequently reconstituted into lethally irradiated young C57BL/6 recipient mice. Shown are the frequencies of peripheral blood (A) CD3e+, (B) CD11b+ and (C) Gr-1+ cells at 2-months post-reconstitution. (D)-(I) These mice were subsequently challenged with a single sub-lethal dose of LPS and peripheral blood immune cells were tracked over time by flow cytometry. Shown are peripheral blood myeloid cells for mice with knockdown of (D) Hoxa9, (E) Ikzf1, (F) Klf4, (G) Klf5, (H) Zbtb4 and (I) Stat3. (J)-(L) These mice were subsequently harvested and the bone marrow compartment was analyzed for the frequency of (J) LT-HSCs, (K) CMPs and (L) CLPs. Data represent at least two independent experiments and are presented as mean ± SEM. * denotes p < 0.05, ** denotes p < 0.01 and *** denotes p < 0.001. P-values was corrected for multiple hypothesis testing by Bonferroni's method.

Of the tested TFs, *Klf5 and Ikzf1* had significant, age-dependent effects. Consistent with the upregulated expression of *Klf5* in mLT-HSCs, we found that knockdown of *Klf5* in aged LT-HSCs resulted in increased lymphoid output (Figure 5A) and decreased myeloid output to levels seen with control young LT-HSCs (Figure 5B,C), whereas its knockdown in young LT-HSCs had no discernable effect compared to controls. Knockdown of *Ikzf1* in young LT-HSCs resulted in decreased lymphoid output (Figure 5A). Interestingly, loss of *Ikzf1* in aged LT-HSCs had no significant effect on lymphoid output, but like knockdown of *Klf5*, it resulted in decreased myeloid output to levels seen with control young LT-HSCs (Figure 5B,C).

We next tested whether these TFs regulate myeloid output of LT-HSCs under conditions of inflammatory stress. To do this, we challenged with LPS the aforementioned mice, each expressing an shRNA of a different TF in the bone marrow compartment (as in Figure 1B-E). The frequency of peripheral blood cells was tracked for 3 weeks after the LPS challenge (Figure 5D-I **and Figure S6B,C**). As expected, mice transplanted with aged LT-HSCs expressing the control vector showed a sustained upregulation of myeloid output over the 3 weeks (Figure 5D-I, solid red lines); whereas mice transplanted with young LT-HSCs expressing the control vector showed only a transient increase in myeloid output 72 hours after the initial challenge, followed by rapid recovery to baseline peripheral blood myeloid frequencies (Figure 5D-I, solid black lines).

In mice transplanted with aged LT-HSCs expressing the *Klf5* shRNA, the response to LPS phenocopied that of mice transplanted with young control LT-HSCs (Figure 5G). These mice responded with a transient increase in myeloid output at 72 hours and rapid recovery to baseline myeloid output, which was lower than that seen with mice transplanted with aged control LT-HSCs (Figure 5G). Similarly, mice transplanted with aged LT-HSCs expressing the *Ikzf1* shRNA had a muted response compared to mice transplanted with aged control LT-HSCs (Figure 5E). These mice sustained myeloid output for 1 week after the initial challenge before returning to their baseline myeloid output, which was equivalent to that seen in mice transplanted with young control LT-HSCs (Figure 5E). Thus, both *Klf5* and *Ikzf1* may play a critical role in regulating inflammatory myelopoiesis in aged LT-HSCs. Mice transplanted with aged LT-HSCs expressing the *Stat3* shRNA also had lower baseline myeloid output than mice transplanted with aged control LT-HSCs, and showed minimal response to LPS (Figure 5I), suggesting *Stat3* may also play an important role in the overall inflammatory response of LT-HSCs.

To identify the cell types responsible for the changes observed after *Ikzf1*, *Stat3* or *Klf5* knockdown in aged HSPCs, we analyzed the bone marrow compartment of all mice 3 months post transplantation. Mice transplanted with aged control cells had higher LT-HSC and lower common lymphoid progenitor (CLP) frequencies compared to mice reconstituted with young control cells (Figure 5J-L, compare black bars to dark red bars). Interestingly, knockdown of either *Klf5*, *Ikzf1* or *Stat3* in transplanted aged LT-HSCs resulted in a decreased frequency of LT-HSCs compared to control transplanted aged LT-HSCs, although the effect of *Klf5* knockdown was only marginally statistically significant (Figure 5J). These knockdowns resulted in LT-HSC frequencies and numbers that were comparable to mice transplanted with young control LT-HSCs (Figure 5J). No significant effect on LT-HSC bone marrow frequency was seen in mice transplanted with young LT-HSCs expressing any of these knockdown constructs (Figure 5J, grey bars).

Some decrease in bone marrow CMP frequency and an increase in CLP frequency was seen in mice transplanted with aged LT-HSCs expressing the *Ikzf1* shRNA when compared with control aged LT-HSCs, however these differences were not statistically significant (Figure 5K,L). This trend may suggest that *Ikzf1* knockdown in aged LT-HSCs phenocopies control young LT-HSCs. No statistically significant changes were observed in the frequencies of other progenitors, including ST-HSCs, MPPs and MEPs when comparing mice transplanted with aged LT-HSCs expressing one of the knockdown constructs compared to aged control LT-HSCs (**Figure S6D-F**). These data therefore suggest that *Klf5*, *Ikzf1* and *Stat3* regulate inflammatory myeloid bias in aged LT-HSCs and may do so by altering the function and frequency of LT-HSCs in the bone marrow compartment.

## Discussion

In this work, we demonstrate that LT-HSCs have a heterogeneous response to inflammatory stimuli that is altered with age. We show that even the most multipotent of HSPCs directly respond *in vitro* to TLR ligands with a potent transcriptional response. Using scRNA-seq, we demonstrate that both the young and aged LT-HSC compartments are comprised of at least two distinct subsets of cells with a defined molecular signature, and with age, the LT-HSC population is enriched for myeloid-biased-like LT-HSCs (mLT-HSCs). Of note, it cannot be verified if the mLT-HSCs identified in this work represent irreversible state or plastic/reversible state within this subpopulation. We posit that an increased proportion of mLT-HSCs in the bone marrow is a key driver of emergency myelopoiesis and further identify several transcription factors that regulate steady state and inflammatory myeloid bias in aged LT-HSCs. Together, our data suggests a revised model (Figure 6) of age-related inflammatory myelopoiesis that highlights important contributions from LT-HSCs, the earliest hematopoietic progenitor.

**Figure 6.**
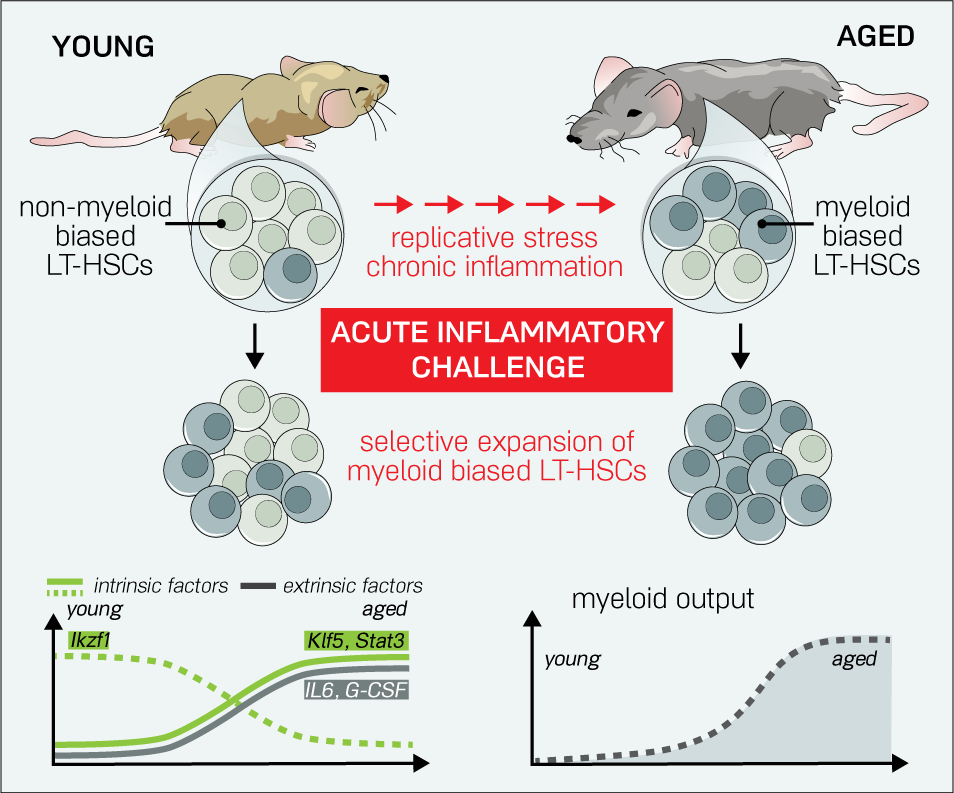
Model of LT-HSC aging and inflammatory myeloid-bias. Shifts in clonal heterogeneity during LT-HSC aging affects the inflammatory response of LT-HSCs. The LT-HSC compartment is comprised of unbiased and myeloid-biased LT-HSCs. With age, the clonal distribution of LT-HSCs shifts towards myeloid-biased variants. During acute inflammatory challenges, myeloid-biased LT-HSCs preferentially expand, leading to increased myeloid output. Several cell-intrinsic factors, including the transcriptional regulators Klf5, Ikzf1 and Stat3 may play a role in establishing a myeloid-biased differentiation program during aging and inflammation. Extrinsic factors, including inflammatory cytokines and growth factors secreted from other cell types may also play a role.

Whereas we found an approximately 30% increase in myeloid cell output in young mice after LPS challenge, this bias is dramatically higher in aged mice, which have a two-fold increase in myeloid output (Figure 6). Interestingly, we found that after inflammatory challenge, aged mice also have a relative increase in LT-HSC frequency in the bone marrow. These effects are likely due to both cell-intrinsic changes in LT-HSCs and changes in the environmental signals present in the bone marrow of aged mice after LPS challenge. We demonstrate a cell-intrinsic component to this differential response between young and aged mice by stimulating LT-HSCs with TLR ligands *in vitro* and transplanting them to lethally irradiated recipients. While it has been demonstrated that TLR stimulation of LT-HSCs induces their proliferation (Zhao et al., 2013), our results suggest that it does not alter their long-term reconstitution potential. Importantly, aged LT-HSCs maintained a memory of the *in vitro* inflammatory challenge and had increased myeloid output 3 months after transplant compared to unstimulated aged LT-HSCs. These data therefore confirms that LT-HSCs directly sense TLR ligands (Nagai et al., 2006), and in response to this, the aged LT-HSC population has an amplified cell-intrinsic myeloid bias. Given that the LT-HSC population is heterogeneous, it is yet to be determined whether inflammation plays a role in either enriching for cells in the mLT-HSC state or in changing the state of nmLT-HSCs to an mLT-HSC-like state.

We show that the various types of HPSCs respond transcriptionally to TLR ligands *in vitro* in a similar way to that seen in BMDCs after LPS stimulation (Ramirez-Carrozzi et al., 2009). Given the different functional roles of HSPCs and mature immune cells, this similarity in transcriptional response is particularly surprising. It has been suggested that, as in mature cell types, inducing the expression of NF-κB with LPS/Pam3csk4 in HSPCs may affect cytokine secretion and proliferation (Zhao et al., 2011, 2014). We observed that the dynamics of expression of NF-κB-driven genes was largely similar between HSPCs and DCs. Thus, the majority of NF-κB-responsive genes appear to be regulated similarly in both HSPCs and mature cells.

Using scRNA-seq, we identified subsets of LT-HSCs with distinct transcriptional responses to inflammatory signals. Previous efforts have successfully identified phenotypic markers for megakaryocyte biased LT-HSC subpopulations (Dykstra et al., 2011; Gekas and Graf, 2013). The gene signature in this study provides functional insight into the basis of myeloid bias in the context of aging and inflammation. Using this myeloid gene signature from stimulated LT-HSCs, we uncovered a subset of mLT-HSC-like cells enriched in the unstimulated aged LT-HSC compartment that have the potential to respond uniquely to acute inflammatory signals. This is consistent with recent results suggesting that there is an epigenetically primed subset of LT-HSCs that is uniquely poised to respond to LPS (Yu et al., 2016). We show herein that such a subset may also be identified using transcriptomic data.

While mLT-HSCs express a gene-signature reflective of myeloid bias, it remains unclear that they preferentially produce myeloid cells in transplant experiments. Since these cells are defined by their transcriptional patterns, and not phenotypic markers, we are unable to perform single-cell transplant experiments to validate their preferential myeloid output with currently available techniques. We do know, however, that the relative distribution of mLT-HSCs as defined by our myeloid-biased gene signature qualitatively reflects the age-related changes in myeloid output we expect to see in LT-HSCs.

Our data supports the model that the increased proportion of these mLT-HSCs with age correlates with an increase in baseline myeloid output in aged mice, which is in turn exacerbated during inflammatory challenge. Accordingly, in the setting of *in vitro* TLR stimulation prior to transplantation (Figure 1F-I), we hypothesize that stimulation of aged LT-HSCs preferentially expands mLT-HSCs or selects cells in an mLT-HSC-like state. This mLT-HSC enrichment then results in a sustained increase in myeloid output for several months post-reconstitution (Figure 1H-I). In the context of physiologic aging, it might be that the accumulation of inflammatory challenges over the lifetime of an animal results in selection and expansion of mLT-HSCs, partially due to direct sensing of these inflammatory signals by these cells. This hypothesis is supported by the fact that chronic inflammatory stimulation of young mice, either by repeated LPS challenge or increased activation of NF-κB, leads to a myeloid-biased output (Esplin et al., 2011; Zhao et al., 2013). However, a direct test of this notion is difficult because even germ-free animals will experience inflammatory events throughout their lifetime. It is therefore possible that there is an intrinsic, inflammation-independent process that drives LT-HSCs towards a myeloid bias over the lifetime of an animal.

Using the inferred myeloid-biased gene signature in our study, we were also able to identify several transcriptional regulators of inflammatory myelopoiesis in aged stimulated mLT-HSCs. Members of the Kruppel-like factor (Klf) family of TFs were among those that were predicted to regulate genes in the myeloid-biased signature and were themselves differentially regulated in mLT-HSCs versus nmLT-HSCs. Among these were *Klf4* and *Klf5*, both of which are required for embryonic stem cell self-renewal (Jiang et al., 2008). *Klf4*, in particular, was one of the key reprogramming factors first used to dedifferentiate somatic cells to induced pluripotent stem cells (Takahashi and Yamanaka, 2016). The enrichment of both of these TFs in mLT-HSCs may therefore play a role in the increased symmetric self-renewal divisions seen in aged LT-HSCs (Geiger et al., 2013; Sudo et al., 2000). Indeed, knockdown of *Klf5* or *Ikzf1* in aged LT-HSCs, but not in young LT-HSCs, results in decreased myeloid output and decreased LT-HSC bone marrow frequency. This suggests that both of these factors may play important roles in regulating LT-HSC myeloid versus lymphoid balance with age. Consistent with these results, it has recently been shown that deficiency of *Klf5* in LT-HSCs leads to decreased bone marrow homing of these cells in transplant experiments and reduced output of myeloid cells, especially neutrophils (Shahrin et al., 2016; Taniguchi Ishikawa et al., 2013). Our results suggest that, in addition to aging, the physiological role of *Klf5* in regulating myeloid output becomes particularly relevant during inflammatory challenge.

The role of *Ikzf1* in LT-HSC function is less well understood. *Ikzf1* is known to regulate early lymphoid differentiation at the level of lymphoid-primed MPPs (LMPPs) and early B and T cell progenitors (Ng et al., 2009; Yoshida et al., 2006). Previous studies have suggested that *Ikzf1* does not play a role in myeloid versus lymphoid lineage commitment of young LT-HSCs (Ng et al., 2009), though some evidence exists that its expression is upregulated in young lymphoid-primed LT-HSCs (Challen et al., 2010). Our results, however, suggest that with age and in the context of inflammation, *Ikzf1* may indeed have a positive role in myeloid fate decisions. Consistent with this, *Ikzf1* has been shown to bind enhancer elements of both myeloid and lymphoid genes in human HSPCs (Novershtern et al., 2011).

Knockdown of *Stat3* in aged LT-HSCs also severely hampered myeloid output after inflammatory challenge. This is consistent with the role of *Stat3* as a major inflammatory TF. In particular, some studies suggest that *Stat3* is induced in response to TLR4 signaling in certain cell types (Kortylewski et al., 2009). Interestingly, complete knockout of *Stat3* in LT-HSCs has been shown to result in a premature aging phenotype (Mantel et al., 2012); our results suggest that partial loss of *Stat3* is not enough to recapitulate this phenotype.

Since decreasing the expression of these TFs can alter the balance of myeloid and lymphoid cells during emergency myelopoiesis, manipulating them or other aspects of the unstimulated or stimulated mLT-HSC programs may provide new therapeutic avenues for re-establishing appropriate lymphoid versus myeloid balance to improve immune function and prevent myeloid leukemias with age.

## Author contributions

M.M., A.M., C.D., M.S.K., A.R. and D.B. designed the experiments. M.M., A.M., M.S.K. and K.L. performed the experiments. C.D. analyzed the sequencing data. A.M., M.M., and D.B. wrote the manuscript, with all authors contributing to writing and providing feedback.

## Acknowledgements

This work was supported by the Sackler Foundation (D.B.), the Howard Hughes Medical Institute (A.R.), the Klaman Cell Observatory at the Broad Institute (A.R.), the Human Frontiers Science Foundation (M.M.), the National Research Service Award CA183220 (A.M), and the UCLA/Caltech Medical Scientist Training Program (A.M), the Canadian Institutes for Health Research (C.G.D), and Charles A. King Trust Postdoctoral Research Fellowship Program, Bank of America, N.A., Co-Trustee and the Simeon J. Fortin Charitable Foundation, Bank of America, N.A. (M.S.K.).

